# Validating marker-less pose estimation with 3D x-ray radiography

**DOI:** 10.1101/2021.06.15.448541

**Authors:** Dalton D. Moore, Jeffrey D. Walker, Jason N. MacLean, Nicholas G. Hatsopoulos

## Abstract

To reveal the neurophysiological underpinnings of natural movement, neural recordings must be paired with accurate tracking of limbs and postures. Here we validate the accuracy of DeepLabCut (DLC) by comparing it to a 3D x-ray video radiography system that tracks markers placed under the skin (XROMM). We record behavioral data simultaneously with XROMM and high-speed video for DLC as marmosets engage in naturalistic foraging and reconstruct three-dimensional kinematics in a shared coordinate system. We find that DLC tracks position and velocity of 12 markers on the forelimb and torso with low median error (0.272 cm and 1.76 cm/s, respectively) corresponding to 2.9% of the full range of marker positions and 5.9% of the range of speeds. For studies that can accept this relatively small degree of error, DLC and similar marker-less pose estimation tools enable the study of more naturalistic, unconstrained behaviors in many fields including non-human primate motor control.

**Summary Statement:** We validate the accuracy of DeepLabCut, a widely used marker-less pose estimation tool, using a marker-based 3D x-ray radiography system (XROMM).

## INTRODUCTION

As the study of motor neuroscience progresses toward an emphasis on naturalistic, unconstrained behavior, kinematics must be captured accurately and efficiently. Past research has relied on marker-based systems that track markers attached to an animal’s skin (such as Vicon, OptiTrack, and PhaseSpace) or surgically implanted radiopaque beads (XROMM; Brainerd et al. 2010). However, these systems are expensive and can be impractical with smaller species like mice or marmosets; they become increasingly impractical if the study focuses on freely-moving behaviors. To solve this problem, multiple groups have developed marker-less pose estimation tools that use deep learning to apply digital markers to recorded video. The most widely used is DeepLabCut (DLC; Mathis et al. 2018), but multiple alternatives exist (Graving et al., 2019; Pereira et al., 2019; Wu et al., 2020). These enable the study of a wider range of behaviors by allowing the subject to move without the disturbance of physical markers. Furthermore, the use of artificial neural networks removes the bottleneck of tracking physical markers semi-automatically; a well-trained, robust network can label video effectively with minimal hands-on time. DLC has been used in a variety of contexts, including tracking eye movements and pupil dilation (Siegle et al., 2019; Steinmetz et al., 2019), hand movements (Sauerbrei et al., 2020), and gross position (Comer et al., 2020) of mice, all with accuracy comparable to human labelers. However, to our knowledge DLC accuracy has not yet been compared to that of marker-based tracking (but see Pouw, Trujillo, and Dixon 2020 for validation in gesture-speech synchrony as compared to marker-based tracking). Such a comparison is crucial to determine whether DLC can reliably track kinematics with accuracy similar to existing marker-based systems and to understand the sources of error that can be ameliorated in future work. XROMM provides a useful basis for comparison, as we have shown that the system tracks radiopaque markers with 0.06 mm precision based on the standard deviation of inter-marker distances during a recording of a calibration specimen (Walker et al., 2020). To this end, we collect simultaneous recordings with XROMM and video (two FLIR Blackfly S cameras at 200 frames per second) as common marmosets engage in naturalistic foraging, then reconstruct three-dimensional reaching kinematics in a shared coordinate system. We find that DLC tracks position and velocity with a median absolute error of 0.272 cm (mean absolute error of 0.344 cm) and 1.76 cm/s (2.50 cm/s), respectively. These errors are 2.9% of the full range of marker positions and 5.9% of the range in speeds. We then discuss sources of error and ways by which this error can be reduced or eliminated.

## METHODS

### Subjects

These experiments were conducted with two common marmosets (*Callithrix jacchus)* (an 8-year old, 356g male and a 7-year old, 418g female). All methods were approved by the Institutional Animal Care and Use Committee of the University of Chicago.

### Data Collection

The two marmosets were placed together in a 1m × 1m × 1m cage with a modular foraging apparatus attached to the top of the cage, as previously described by Walker et al. (2020). The marmosets were allowed to forage voluntarily throughout recording sessions that lasted 1-2 hours. Individual trials were triggered manually with a foot pedal by the experimenters when the marmosets appeared ready to initiate a reach. The manual trigger initiated synchronized video collection by the XROMM system (Brainerd et al., 2010) and two visible light cameras, each described in further detail below. We retained all trials that captured right-handed reaches.

### XROMM

Bi-planar X-ray sources and image intensifiers (90kV, 25mA at 200 fps) were used to track the 3D position of radiopaque tantalum beads (0.5-1 mm, Bal-tec) placed subcutaneously in the arm, hand, and torso. Details of bead implants can be found in Walker et al. (2020), in which the authors also report estimating XROMM marker tracking precision of 0.06 mm. Positions of 13 beads were tracked using a semi-automated process in XMALab (Knorlein et al., 2016). One bead implanted in the torso was ignored for analysis later to match the 12 labels in DLC.

### DeepLabCut

Two high-speed cameras (FLIR Blackfly S, 200 fps, 1440×1080 resolution) were used to record video for analysis by DLC. The cameras were positioned to optimize visibility of the right upper limb during reaching behavior in the foraging apparatus and to minimize occlusions, while avoiding the path between the X-ray sources and image intensifiers (Fig. 1A). The cameras were triggered to record continuous images between the onset and offset of the manual XROMM trigger, with series of images later converted to video for DLC processing. We labeled 12 body parts in DLC – three labels on each of the torso, upper arm, forearm, and hand (Fig. 1B). We used DLC 2.0.7 with in-house modifications to produce epipolar lines in image frames that were matched between the two cameras (Fig. 1C), which significantly improved human labeling accuracy. This modification has since been added as a command line feature in the DLC package. Aside from this and related changes to the standard DLC process, we followed the steps outlined in Nath et al. (2019).

**Figure 1:**
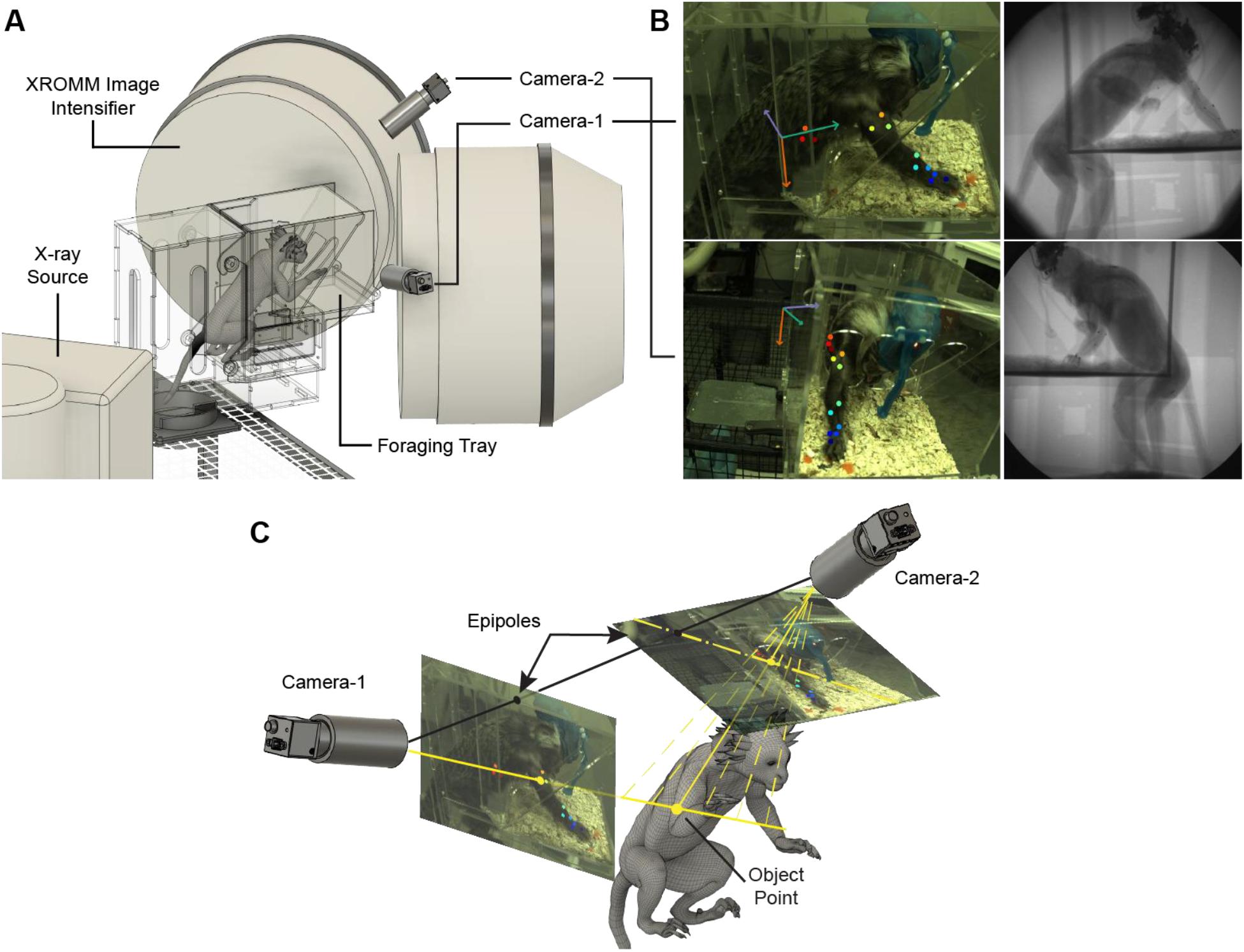
Recording apparatus, markers, and labeling with epipolar lines. A) Marmosets use their upper limb to forage. Blackfly S cameras and XROMM record foraging behavior simultaneously. B) *Left:* Videos from camera-1 (top) and camera-2 (bottom). Markers are applied by DLC. The coordinate system generated by camera calibration is shown in green (x), orange (y), and purple (z). *Right:* corresponding frames from XROMM. Markers are tantalum beads implanted subcutaneously, seen as small black dots and tracked offline with XMALab. Coordinate basis (not shown) corresponds to that of DLC due to bimodal calibration. C) Epipolar lines improve labeling accuracy. A vector projects from camera-1 through the applied label into 3D space, where it intersects with possible vectors from camera-2 (a subset is shown in dotted yellow). The epipolar line (dot-dash) passes through the epipole in camera-2 and each of the points at which a vector intersects the image plane. A correct label applied along the epipolar line produces accurate triangulation to the object point (correct labels and camera-2 vector shown in solid yellow).

### Calibration

A custom calibration device was built to allow for calibration in both recording domains (Knorlein et al. 2016; instruction manual for small lego cube is located in the XMALab BitBucket). The device was constructed from LEGO bricks and Sugru putty, with a three-dimensional grid of steel beads within the structure and a two-dimensional grid of white circles on one face of the cube. Calibration of x-ray images was computed in XMALab and calibration of visible light images was computed with custom code using OpenCV. This integrated calibration device ensures that DLC and XROMM produced tracked trajectories in a common 3D coordinate system.

### Training network and experimental design

We trained two separate networks to label marmoset body parts – one for each marmoset. Marmoset TY produced four useful reaching events containing 1-3 reaches each and marmoset PT produced 13 reaching events. We extracted 100 total frames (50/camera) across the four events for marmoset TY and 254 frames (127/camera) across seven events for marmoset PT, although we later removed one these (and its eight pairs of images) and one of the unlabeled events from the data because they each contained a single reach with so many occlusions that they could not be adequately tracked without extensive refinement. After preprocessing (discussed below), our dataset consisted of 5721 frames in the test set and 169 frames in the train set. All 169 training frames came from four TY events and six PT events, while the test frames came from those same events as well as five PT events that were completely held-out from the training set.

### Label refinement

Both networks labeled reaching events well in the case that the reach trajectory was represented in the training data. As documented by the DLC authors, the networks failed in sections of reaches that were similar to poorly labeled training images or sections of reaches completely unrepresented in the training set. After the first iteration of video analysis was completed and evaluated, we used the DLC refinement tools to extract outlier frames in poorly labeled videos and improve the training set before retraining the network. We went through the refinement process one time.

### Trajectory processing

Because we focused on labeling periods of active reaching, large portions of video that capture the marmoset chewing or otherwise disengaged from the foraging task were poorly labeled by the DLC networks. In order to isolate reaching movements, we first removed all frames for which the label’s DLC likelihood score was below 0.05. We then removed all frames for which the lateral hand label was well behind the foraging partition and retained the rest of the frames. The remaining frames correspond to reaching movements within the foraging arena and brief periods of rest preceding, between, and after reaching movements. DLC trajectories were smoothed using a 3^rd^ order Savitsky-Golay filter with a 31-sample sliding window. Finally, we found that there was a brief delay ranging from 0 to 10 frames between pedal-triggered onset of the XROMM event and the corresponding pedal-triggered TTL pulse initiating the start of the event for the FLIR cameras (and for the pulse ending the event). To adjust for the timing difference, we iterated over a range of possible sample shifts separately for each event to find the shift that minimized the mean absolute error between DLC and XROMM trajectories. We visually inspected each trajectory after the adjustment to ensure the shift was qualitatively accurate.

### Evaluation of DLC Accuracy

We computed the median and mean absolute error between the position and velocity traces found using DLC and the matched XROMM trajectories for all body parts across all reaching events. Body parts were ignored for instances in which they were obscured from view by the visible light cameras for an extended period as a result of the second marmoset entering the camera view mid-reach. This applied to the torso markers in one event and all but the hand markers in another event.

### Statistical Tests

Since the error distributions are right-skewed with long tails of large errors, we use the median error to describe the center of each distribution and the Mann-Whitney U-Test to assess statistical significance. The P-values computed with this method are artificially low due to the large sample size (e.g. 17,671 samples for the three hand markers and 14,986 samples for the three torso markers), so we report the correlation effect size defined by the rank-biserial correlation to describe statistical differences between distributions. According to convention, we consider r < 0.20 to be a negligible effect (Cohen, 1992).

### Normalized Error and Fraction of Variance Accounted For

To compute normalized error, we divided the position and velocity errors by the maximum range of motion and maximum speed for each marker across the dataset. To compute the fraction of variance accounted for, we used the following equation:

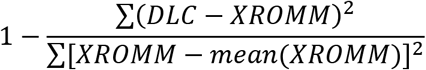

which normalizes the sum of squared DLC error by the XROMM variance and subtracts from one.

## RESULTS

### Position and Velocity Error – Tracking Examples

Qualitatively, DLC and XROMM-tracked forelimb trajectories were nearly identical for most time points and reaches (Fig. 2A-C). The 3D view shows DLC tracking the overall motion and most fine details accurately, while missing some brief position changes tracked by XROMM near the start of the reach (Fig. 2A). Breaking out x-y-z components demonstrates minimal position and velocity error (Fig. 2B,C), with median error of 0.238 cm and 2.25 cm/s (mean error of 0.237 and 2.70 cm/s) for this event.

**Figure 2:**
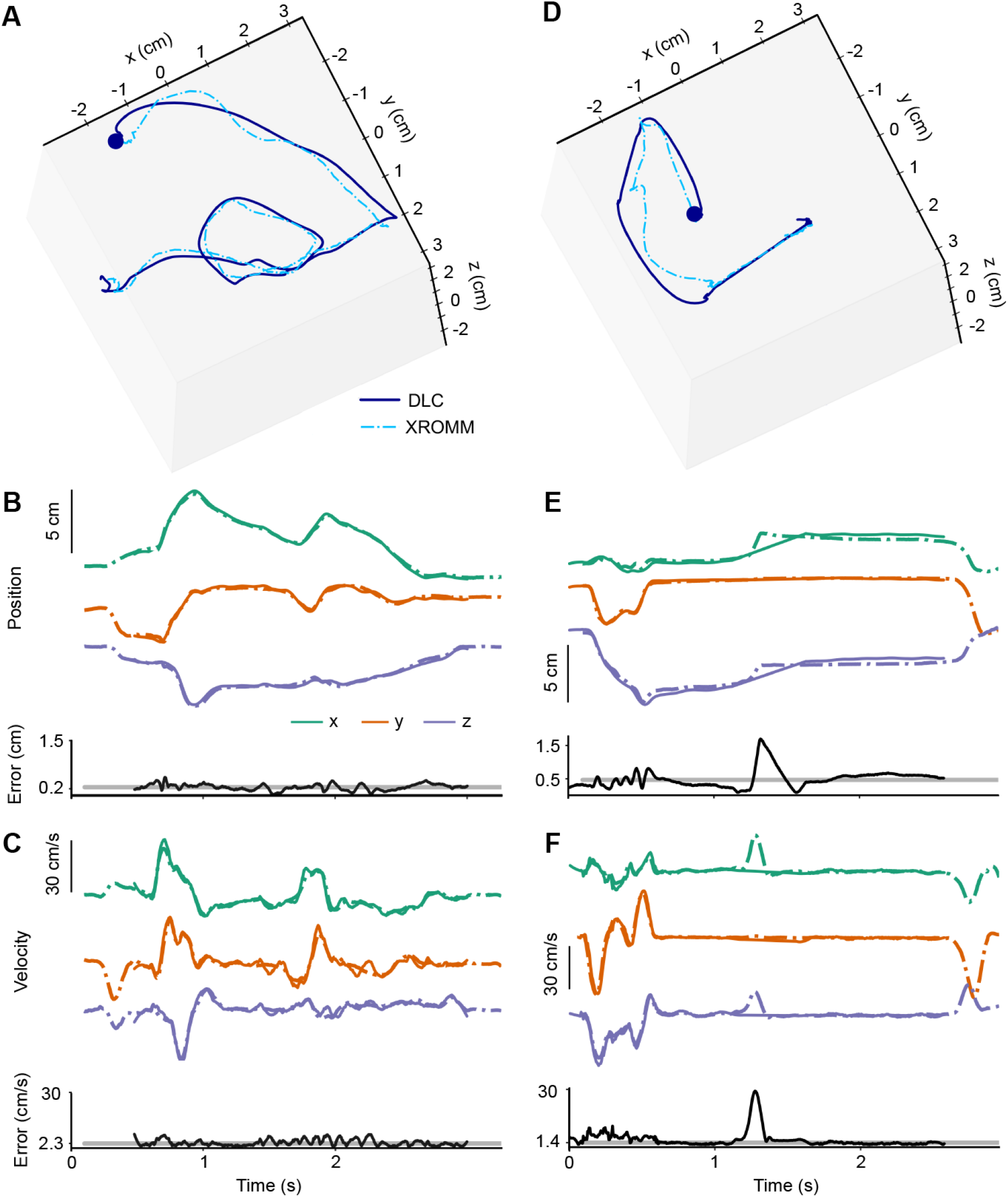
Tracking position and velocity with DLC and XROMM. A-C) Example of accurate DLC tracking. A) 3D position of distal hand marker with DLC (dark blue, solid) and XROMM (light blue, dot-dash). The plot is rotated to approximate the DLC camera-1 angle. B) Detailed position. *Top:* kinematics split into x-y-z coordinates colored to match the axes in Fig. 1B, with DLC in solid and XROMM in dot-dash. XROMM extends before and after DLC because we filtered out DLC when the hand is well behind the foraging partition. *Bottom:* Error at each time (black) overlaid on median error for the event (gray). C) Velocity for same event. D – F) Example of the errors produced by DLC. The marker movement missed by DLC at ~1.25s came from removing a marker jump during a movement and replacing it with smoothed interpolation.

Some reaches were not tracked with the same accuracy, with large errors falling into a few categories. One category is exemplified in Fig. 2D-F, in which a marker jump that occurred around 1.25s was removed during DLC processing and replaced with a linear interpolation (Fig. 2E,F). This results in a spike of position and velocity error and correspondingly high average errors (median of 0.464 cm and mean of 0.488 cm for position, median of 1.37 cm/s and mean of 2.88 cm/s for velocity). A second category of errors occurred when markers moved rapidly away from the correct location over a few frames at the beginning or end of tracking. The majority of these large errors occurred because the training dataset did not contain frames that were similar enough to the problematic reaching periods (see Discussion). Finally, some of the less extreme errors above the median occurred due to human labeling inconsistency, particularly when visual landmarks were unclear or changed due to skin deformation.

### Position and Velocity Error – Aggregate Results

We found that DLC tracked position and velocity with median error of 0.272 cm (mean of 0.344 cm) and 1.76 cm/s (2.49 cm/s), respectively (Fig. 3A,B, right). Position and velocity error distributions were right-skewed with a long tail of large errors, so we focused on median error as the measure of accuracy. We found that position errors for the forearm (median of 0.381 cm) and torso (0.328 cm) were significantly larger than errors for hand (0.217 cm) and upper arm (0.215 cm) markers (Mann Whitney U-test; r_forearm-hand_ = 0.42, r_forearm-upperArm_ = 0.46, r_torso-hand_ = 0.34, r_torso-upperArm_ = 0.38). Velocity errors varied between body segments to a lesser degree – the only modest difference was higher error for the torso than the upper arm (medians of 1.93 cm/s and 1.47 cm/s, r = 0.23). We note that the forearm and torso were more difficult to label manually than the hand and upper arm. As such, we attribute variation in tracking quality across body segments to human labeling inconsistency rather than any issues with DLC itself; we suggest taking the hand and upper arm errors as more indicative of DLC performance.

**Figure 3:**
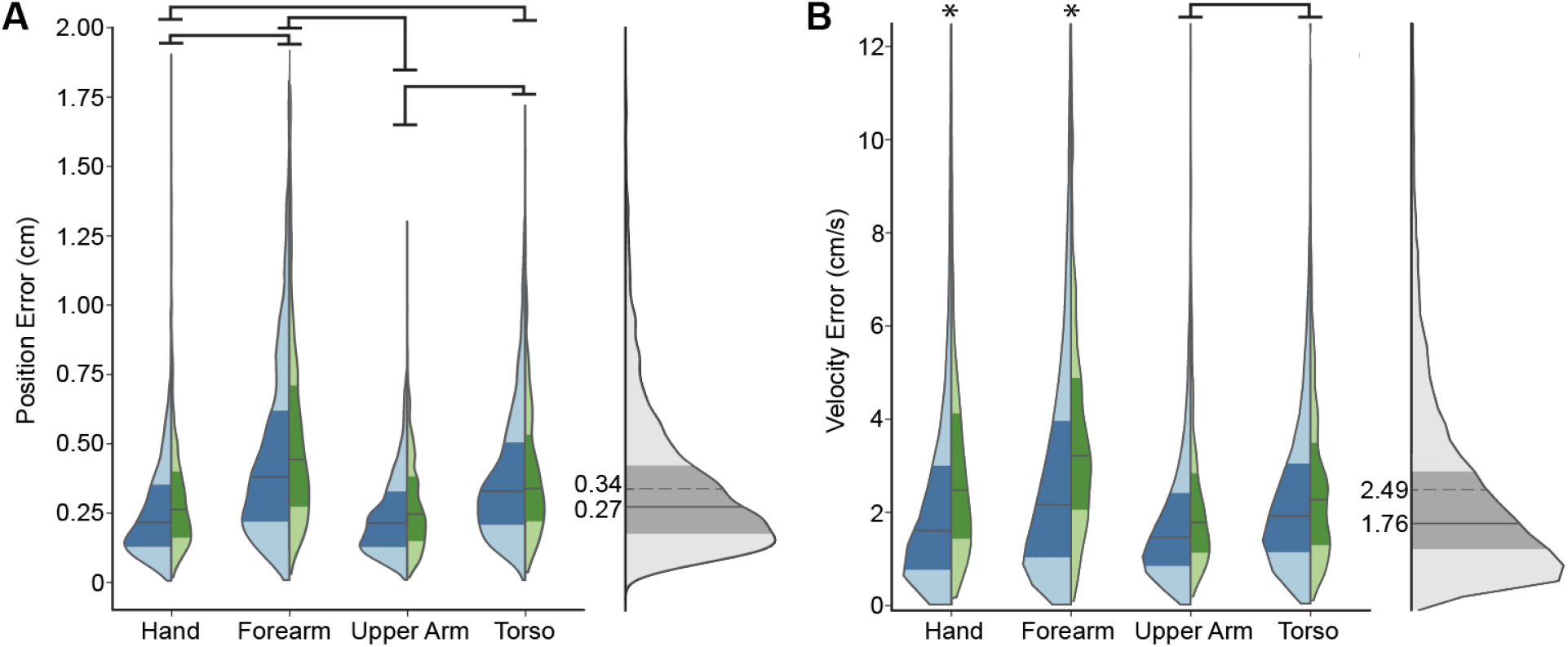
Error distributions. A) Position error distributions. *Left:* violin plots of absolute error separated by body segment. Each segment contains three DLC-XROMM marker pairs. Errors for 169 training frames are in green, with 5,721 test frames in blue. Median and interquartile range are shown by a gray line and shading. Significant effects with r>0.20 are indicated by bars for segments and asterisks for test-train. *Right:* distribution of pooled errors using the same conventions for median and interquartile range, and mean error at the dotted line. B) Velocity error distributions.

To provide context, we normalized position and velocity errors by the maximum range of position and speed for each marker. We found that the median error across events and markers was just 2.9% of position range and 5.9% of the speed range. The hand markers, for which visual landmarks were most clear, had median errors equivalent to 1.9% and 3.2% of position and speed ranges. Viewed another way, we find that DLC accounts for 94.1% of the XROMM position variance and 81.2% of velocity variance for all data points, or 96.5% and 89.6%, respectively, when we removed errors above the 0.975 quantile to focus on well-labeled data.

We separated the 169 datapoints that correspond to image frames labeled by hand (train) from the rest of the data which were labeled by the network only (test) and report the median error across three markers on each body segment (Fig. 3A,B, left). Median position error was 0.32 cm for train and 0.27 cm for test, but this effect was negligible (r = 0.11). Velocity train errors were significantly greater (median = 2.40cm/s) than test (1.75 cm/s) with r = 0.22. The hand (2.48 cm/s to 1.61 cm/s, r = 0.27) and forearm (3.21 cm/s to 2.17 cm/s, r = 0.29). drove this result, while the effect was negligible for the upper arm and torso.

### Inter-Marker Distances

We estimated the precision of tracking by the standard deviation of inter-marker distances within each body segment, with the expectation that inter-marker distances should be near constant aside from soft tissue deformation. We report precision of 0.21 cm for the hand and forearm, 0.22 cm for the upper arm, and 0.37 cm for the torso. This is of similar magnitude to the position error and results from variability in human labeling and soft tissue deformation.

## DISCUSSION

We have shown that DLC tracks movements of marmoset forelimb and torso accurately in comparison to marker-based tracking with XROMM. The median errors of 0.272 cm and 1.76 cm/s (2.9% and 5.9% of kinematic range) are sufficient for many purposes, and there is reason to treat this as an upper error bound for well-trained DLC networks. These errors should provide an expectation of the accuracy that can be attained with a simple recording setup and relatively little computer vision experience, but which could be improved with targeted adjustments.

### Assessment of Errors and Limitations

We present these results as an upper bound on DLC error due to some limitations in the dataset. The first is the small number of events due to difficulties associated with the XROMM; the marmosets were uncomfortable in the space and therefore less interested in foraging and it was difficult to manually trigger data collection at the onset of voluntary movements. The second limitation is the steep angle recorded with DLC camera-2 (Fig. 1), which would ideally have been placed in the center of the image intensifier rather than mounted on top – this made identifying visual landmarks difficult, particularly on the torso where only a fraction of potential landmarks was visible in most frames. Finally, the useful landmarks on the forearm did not align well with estimated XROMM bead positions on the DLC images and were difficult to locate when the marmoset made short reaches to the left corner of the foraging tray. Labeling of the hand and upper arm was not as affected by the camera angles and ambiguous landmarks, which explains the lower error for these body segments.

DLC performance is dependent on the quality and breadth of training data - movements that are sparsely or un-represented in training data are unlikely to be well-tracked by the network. Tracking errors such as in Fig. 2D-F can be reduced by labeling a comprehensive dataset from the outset or by rigorous relabeling of problematic video frames using the DLC toolbox.

### Improving Accuracy

DLC tracking can be made more accurate by improving data quality, improving human labeling performance, and utilizing processing tools within and outside of DLC. Image quality is the fundamental limiting factor to DLC accuracy – resolution affects the number of pixels per unit distance, while resolution, visibility, contrast, and brightness constrain human labeling consistency. We suggest recording at the highest feasible resolution and using more than two cameras to improve coverage and reduce occlusions, as well as increase the benefit of 3D processing tools discussed below. Additionally, lighting equipment and consideration of occlusions and visible landmarks on the animal can improve human labeling.

After collecting high-quality data, there are tools available to improve tracking accuracy. Most proximally, DLC is open-source and actively maintained, with frequent implementation of new features. We have contributed one such feature by incorporating epipolar lines to improve labeling precision across cameras for 3D projects (Fig. 1C). Epipolar lines simplify the identification of the same landmark in different camera angles and are especially helpful when landmarks are ambiguous. At time of publication, this has been implemented for pairs of cameras but could be extended such that epipolar lines from two cameras provide a triangulated point in subsequent views.

Additional tools operate as add-ons to base DLC, including anipose (Karashchuk et al., 2020), pose3d (Sheshadri et al., 2020), OpenMonkeyStudio (Bala et al., 2020), and a new development that uses 3D modeling to create synthetic training data (Bolaños et al., 2021). Anipose and pose3d facilitate 3D pose estimation with more than 2 cameras and standardized filtering options. Anipose provides an especially useful 3D-filtering option that improves performance over the standard process of filtering in 2D before triangulation to 3D coordinates. OpenMonkeyStudio and the Bolaños et al. (2021) method are data augmentation tools – the former uses labels on a subset of 62 camera views to produce triangulated labels on remaining views, while the latter animates a 3D model to render photo-realistic images and labels as synthetic training data for DLC.

### Applicability to other animal pose estimation tools

Although DLC is the most widely used marker-less pose estimation tool, there are a few alternatives. LEAP (Pereira et al., 2019) and DeepPoseKit (Graving et al., 2019) use different network architectures to reduce training and inference time. Deep Graph Pose (DPG; Wu et al. 2020) uses the same architecture as DLC but implements temporal and spatial constraints to reduce the necessary training set size and eliminate marker jumps. However, the authors do not compare performance for the frames without marker jumps, which don’t seem to differ from DLC in example trials they present. This suggests that DPG reduces the long tail of high-error frames, but it is unclear whether it improves accuracy for well-tracked frames. Thus, we expect DLC errors reported here to be representative for existing tools, at least for the subset of well-tracked frames.

### Future Work

We plan to extend DLC outputs to reconstruct joint angular motion about the shoulder, elbow, and wrist in a joint coordinate system following Walker et al. (2020), by adapting the workflow described by Brainerd et al. (2010). However, dealing with inter-marker distance jitter and estimating the position of DLC labels on 3D-rendered CT-scan models has proven challenging. We hope to address these problems and decrease “human” labeling inconsistency by generating synthetic training data with a 3D marmoset model (similar to the mouse model described by Bolaños et al. 2021) and applying DLC outputs with this marmoset model to constrain the inverse kinematics computations of joint angular motion.

## ACKNOWLEDGEMENTS

We thank Marina Sundiang for early help setting up data collection and for contributions to animal care and training; J.D. Laurence-Chasen for help building the Lego calibration object; the XROMM Technology Development Project for the XROMM; Ben Knorlein for XMALab and assistance locating pertinent code for rigid body transformations in the XMALab code; and Alexander Mathis and Mackenzie Mathis for DeepLabCut. This is University of Chicago XROMM Facility Publication 8.

## AUTHOR CONTRIBUTIONS

D.D.M, J.D.W., J.N.M. and N.G.H. developed the initial concept of validating DLC with XROMM. D.D.M. and J.D.W. set up and optimized cameras for DLC and collected all data. D.D.M. analyzed all the data and wrote all the code, with significant help and input from J.D.W.; D.D.M. wrote the manuscript and all authors read the manuscript, provided critical comments for revision and approved the final version. J.N.M. and N.G.H. provided guidance at each phase of the process. D.D.M., J.D.W., J.N.M. and N.G.H. acquired funding.

## DATA AVAILABILITY

Data available upon request. Code available at https://github.com/hatsopoulos-lab/marmoset-dlc_xromm_validation.

